# A Comprehensive epidemiological and molecular study of gastrointestinal helminths of Companion Animals in Northeastern Bangladesh: A neglected zoonotic threat

**DOI:** 10.64898/2026.02.05.704122

**Authors:** Tilak Chandra Nath, Jannatul Nyema, Radwan Raquib, Tarek Siddiki, Jarin Tasnim, Zakia Sultana Prity, Mohiuddin Tarek, Mandira Mukutmoni, Kazi Mehetazul Islam, Sultan Ahmed

## Abstract

**Background:** Gastrointestinal helminths of companion animals are neglected sources of zoonotic infection in low and middle-income countries. In Bangladesh, close human-animal contact and large free-roaming dog and cat populations may facilitate parasite transmission, yet region-specific data remain limited. This study assessed the prevalence, species diversity, and zoonotic potential of gastrointestinal helminths in companion animals in northeastern Bangladesh.

**Methods:** A cross-sectional study was conducted between January and December 2025 across urban and rural areas of the Sylhet Division. Fecal samples from 900 animals (600 dogs and 300 cats; owned and stray) were examined using standard coproscopic techniques. Molecular confirmation of selected positive samples was performed using PCR targeting ITS-2, 18S rRNA, and mitochondrial cox1 genes, followed by sequencing. Risk factors associated with infection were evaluated using multivariable logistic regression.

**Results:** Overall, 45.9% (95% CI: 42.6–49.2) of animals were infected with at least one gastrointestinal helminth, with mixed infections detected in 18.4%. Prevalence was similar in dogs (45.7%) and cats (46.3%) but significantly higher in stray animals (65.7%) than in owned animals (36.6%). Predominant zoonotic helminths included *Ancylostoma spp*., *Toxocara canis, Toxocara cati, Dipylidium caninum*, and Taenia/*Echinococcus spp*. Molecular analysis confirmed 93% of morphologically identified infections and revealed high genetic similarity to zoonotic reference strains. Stray status, lack of deworming, young age, and outdoor roaming were significant risk factors for infection (p < 0.05).

**Conclusions:** Companion dogs and cats in northeastern Bangladesh harbor a high burden of zoonotic gastrointestinal helminths and represent important reservoirs for human exposure. Strengthening One Health–based surveillance, routine deworming, and stray animal management is essential to reduce zoonotic transmission.

## Introduction

Gastrointestinal helminth infections in companion animals, particularly dogs and cats, represent a persistent yet underrecognized public health concern in many low and middle-income countries (LMICs), including Bangladesh. High human population density, inadequate sanitation infrastructure, and close human-animal contact create favorable conditions for the transmission of zoonotic diseases [1,2]. Infected companion animals often suffer from chronic health consequences such as malnutrition, anaemia, impaired growth, and reduced immunity, while simultaneously serving as reservoirs for parasites of significant zoonotic importance [3]. Despite their dual impact on animal and human health, gastrointestinal helminths of companion animals remain largely neglected in national disease surveillance and control programs in Bangladesh.

Sylhet district in northeastern Bangladesh is characterized by a humid subtropical climate, rapid and unplanned urbanization, and a substantial population of free-roaming and semi-domesticated dogs and cats. These ecological and socio-demographic factors collectively increase the risk of environmental contamination with infective helminth stages and facilitate zoonotic transmission through soil, water, and direct contact [4,5]. Children are particularly vulnerable due to frequent soil exposure and suboptimal hygiene practices. Among the most clinically significant zoonotic helminths, *Toxocara canis* and *Toxocara cati* cause visceral and ocular larva migrans in humans, leading to severe morbidity and long-term sequelae [6]. Hookworms such as *Ancylostoma caninum* are responsible for cutaneous larva migrans, while *Dipylidium caninum* and *Echinococcus spp*. pose additional risks, including cystic echinococcosis with potentially fatal outcomes [7]. Although several studies have documented helminth infections in livestock in Bangladesh, systematic investigations focusing on companion animals remain limited, particularly at the regional level [8]. Existing data suggest a high burden of infection, yet the true prevalence, species composition, and zoonotic potential of gastrointestinal helminths in dogs and cats are poorly defined. Conventional coproscopic methods are widely used in field settings; however, their limited sensitivity and inability to reliably distinguish morphologically similar species restrict accurate risk assessment. The integration of molecular diagnostic tools, such as polymerase chain reaction (PCR), is therefore essential to achieve species-level identification and improve understanding of transmission dynamics [9].

Recent national-scale investigations have emphasized the need for risk-based surveillance and a One Health approach to address parasitic zoonoses in Bangladesh [10]. Nevertheless, region-specific evidence from northeastern Bangladesh, particularly Sylhet, remains scarce, creating a critical knowledge gap that hinders effective policy formulation and targeted intervention strategies. Addressing this gap is especially important given the region’s unique ecological conditions and intense human-animal-environment interactions.

Against this backdrop, the present study provides the first comprehensive epidemiological assessment of gastrointestinal helminths in companion dogs and cats across urban and rural settings in Sylhet, northeastern Bangladesh. By integrating coproscopic examination with PCR-based molecular confirmation, this study aims to determine parasite prevalence, species diversity, and associated risk factors. The findings are intended to elucidate the zoonotic potential of gastrointestinal helminths in companion animals, identify high-risk transmission settings, and generate evidence to support targeted control strategies within a One Health framework.

## Materials and Methods

### Study Design and Geographical Setting

A cross-sectional epidemiological study was conducted between January and December 2025 across urban and rural ecological settings of Sylhet, northeastern Bangladesh. Sylhet lies within a humid subtropical climatic zone characterized by high annual rainfall (∼4,000 mm), elevated humidity, and temperatures favourable for the development and survival of helminth eggs and larvae. Urban sampling was conducted in Sylhet Sadar, a densely populated metropolitan area (∼15,000 persons/km^2^) with extensive concrete infrastructure, limited green space, and a high density of owned and free-roaming companion animals. Rural sampling encompassed five sub-districts (upazilla) representing diverse ecological and socio-environmental conditions: Balaganj (24°55′N, 91°52′E), dominated by tea plantations and wetland ecosystems; Golapganj (24°50′N, 92°00′E), characterized by riverine habitats; Biswanath (24°41′N, 91°47′E), featuring mixed crop-livestock farming systems; Jaintiapur (25°07′N, 92°08′E), located in a hilly border region; and Fenchuganj (24°40′N, 91°55′E), representing industrial-agricultural interfaces. These locations were purposively selected to capture variability in land use, climate, sanitation, and human-animal contact patterns influencing helminth transmission [11].

### Ethical Approval and Animal Welfare

The study protocol was reviewed and approved by the Animal Research Ethics Committee of Sylhet Agricultural University, Bangladesh. Animal handling was performed by trained veterinarians to minimize stress and discomfort. Oral informed consent was obtained from pet owners before sample collection after providing a clear explanation of the study objectives, procedures, and potential benefits.

### Study Population and Sampling Strategy

The study population comprised domestic and free-roaming companion dogs and cats. A total of 900 animals (600 dogs and 300 cats) were enrolled using a stratified random sampling strategy based on study sites and ownership status (owned vs. stray). Of these, 600 were owned animals (400 dogs and 200 cats) recruited from households and veterinary clinics, while 300 were free-roaming animals (200 dogs and 100 cats) sampled opportunistically with community assistance. Sample size was calculated following the method described by Thrusfield [12] for prevalence studies, assuming an expected prevalence of 42% with 95% confidence level, and 5% margin of error, based on previous studies conducted in Bangladesh [13]. The final sample size was increased to enhance statistical power and account for potential sample loss.

### Sample Collection and Preservation

Fresh fecal samples were collected immediately after defecation, either directly from the environment for free-roaming animals or during routine handling of owned animals. A minimum of 5 g of feces was collected per animal and divided into two sterile, labeled containers: one aliquot without preservative for coproscopic examination and a second aliquot preserved in 70% ethanol for molecular analysis. Samples were transported in insulated containers to the Parasitology Laboratory of Sylhet Agricultural University and processed within 24 hours of collection.

### Coproscopic Examination

All fecal samples were examined using a combination of qualitative and concentration techniques to maximize diagnostic sensitivity. Formalin-Ether Concentration Technique, Sheather’s sugar flotation method and Modified Baermann technique for larval detection were employed [14]. Parasite eggs and larvae were identified based on standard morphological criteria.

### Molecular Identification

Molecular characterization was performed on 100 coproscopically positive samples, stratified by host species, parasite group, and geographic location. Genomic DNA was extracted using the QIAamp DNA Stool Mini Kit (Qiagen, Germany) following the manufacturer’s protocol with minor modifications to enhance DNA yield. Polymerase chain reaction (PCR) assays targeting different helminth groups were conducted using established primer sets using both mitochondrial cytochrome c oxidase subunit I (*cox*1) and 18S rRNA ITS region primers. Each PCR reaction contained 2 μL of template DNA, 12.5 μL of DreamTaq Green PCR Master Mix (Thermo Fisher Scientific), 1 μL of each primer (10 μM), and nuclease-free water to a final volume of 25 μL. Thermocycling conditions consisted of an initial denaturation at 95°C for 5 min, followed by 35 cycles of denaturation at 95°C for 30 s, annealing at 55°C for 45 s, and extension at 72°C for 1 min, with a final extension at 72°C for 7 min. PCR products were visualized on 1.5% agarose gels using a Gel Doc XR+ imaging system (Bio-Rad). Amplified products were purified using the DokDo-Prep PCR Purification Kit (ELPIS Biotech, South Korea) and bidirectionally sequenced by BDGenome (Bangladesh). Sequence editing and alignment were performed using ClustalW implemented in Geneious v9 software. Species identification was confirmed by comparison with reference sequences deposited in GenBank using BLAST.

### Questionnaire Survey and Statistical Analysis

A structured questionnaire survey was conducted with 200 pet owners and 50 practicing veterinarians to collect data on animal demographics, husbandry and management practices, deworming history, environmental exposure, and perceived zoonotic health risks. Data were entered into Microsoft Excel and analyzed using R software version 4.3.0. Statistical analyses were initially performed to screen potential risk factors. Variables with a p-value <0.20 were subsequently included in multivariable logistic regression models using backward stepwise elimination. Adjusted odds ratios (AOR) with 95% confidence intervals (CI) were calculated, and statistical significance was set at p < 0.05.

## RESULTS

### Overall prevalence of gastrointestinal helminths in dogs and cats

A total of 900 fecal samples obtained from companion dogs and cats in northeastern Bangladesh were examined for gastrointestinal helminths. Overall, 413 samples were positive for at least one helminth species, yielding an overall prevalence of 45.9% (Table 1, Fig 1). Mixed infections involving two or more helminth taxa were detected in 18.4% (76/413) of infected animals. The prevalence was comparable between dogs and cats.

**Table 1.** Overall prevalence of gastrointestinal helminths in dogs and cats.

| Host species | No. examined | No. positive | Prevalence (%) | 95% CI |
| --- | --- | --- | --- | --- |
| Dogs | 600 | 274 | 45.7 | 41.7-49.7 |
| Cats | 300 | 139 | 46.3 | 40.7-52.0 |
| Total | 900 | 413 | 45.9 | 42.6-49.2 |

**Table 2.** Species-specific prevalence of gastrointestinal helminths in dogs.

| Helminth species | No. positive | Prevalence (%) | 95% CI |
| --- | --- | --- | --- |
| Hookworms | 169 | 28.2 | 24.6-31.8 |
| *Toxocara canis* | 88 | 14.7 | 11.9-17.5 |
| *Trichuris vulpis* | 59 | 9.8 | 7.4-12.2 |
| *Dipylidium caninum* | 56 | 9.3 | 7.0-11.7 |
| *Taenia/Echinococcus* spp. | 21 | 3.5 | 2.0-4.9 |
| *Diphylobothrium* spp. | 3 | 0.5 | 0.0-1.1 |
| Larvae (unspecified) | 39 | 6.5 | 4.5-8.4 |

**Table 3.**
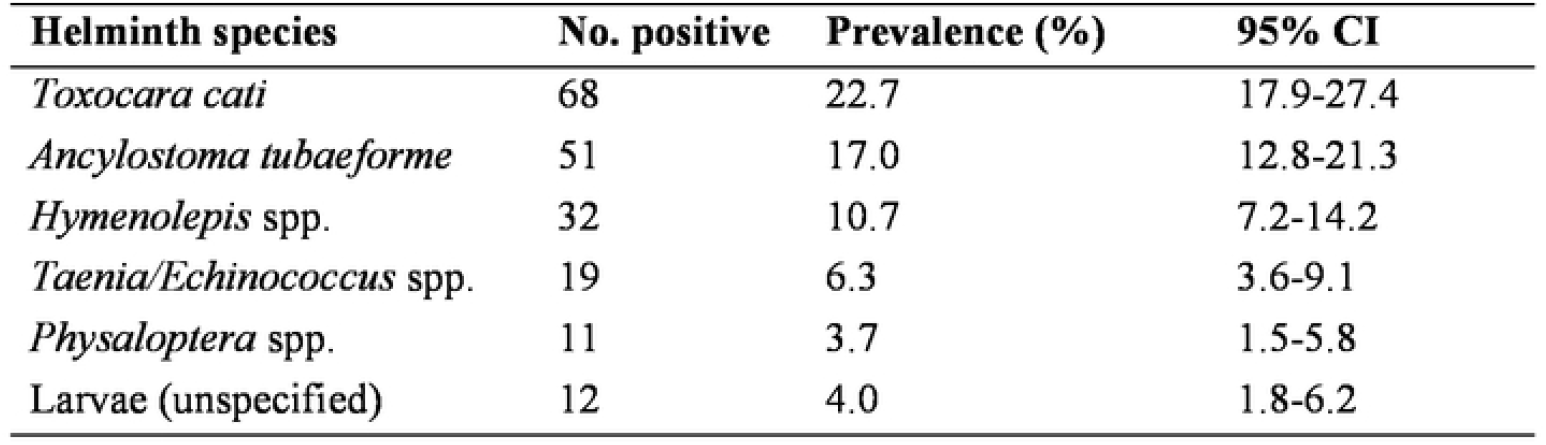
Species-specific prevalence of gastrointestinal helminths in cats.

**Table 4.** Prevalence of gastrointestinal helminths in pet-owned animals.

| Animal type | No. examined | No. positive | Prevalence (%) | 95% CI |
| --- | --- | --- | --- | --- |
| Dogs | 400 | 142 | 35.5 | 31.2-39.6 |
| Cats | 200 | 78 | 39.0 | 29.4-48.6 |
| Total | 600 | 220 | 36.6 | 32.2-39.8 |

**Table 5.** Prevalence of gastrointestinal helminths in stray animals.

| Animal type | No. examined | No. positive | Prevalence (%) | 95% CI |
| --- | --- | --- | --- | --- |
| Dogs | 200 | 132 | 66.0 | 61.6-69.4 |
| Cats | 100 | 61 | 61.0 | 57.4-65.7 |
| Total | 300 | 193 | 64.3 | 60.3-71.0 |

**Table 6.**
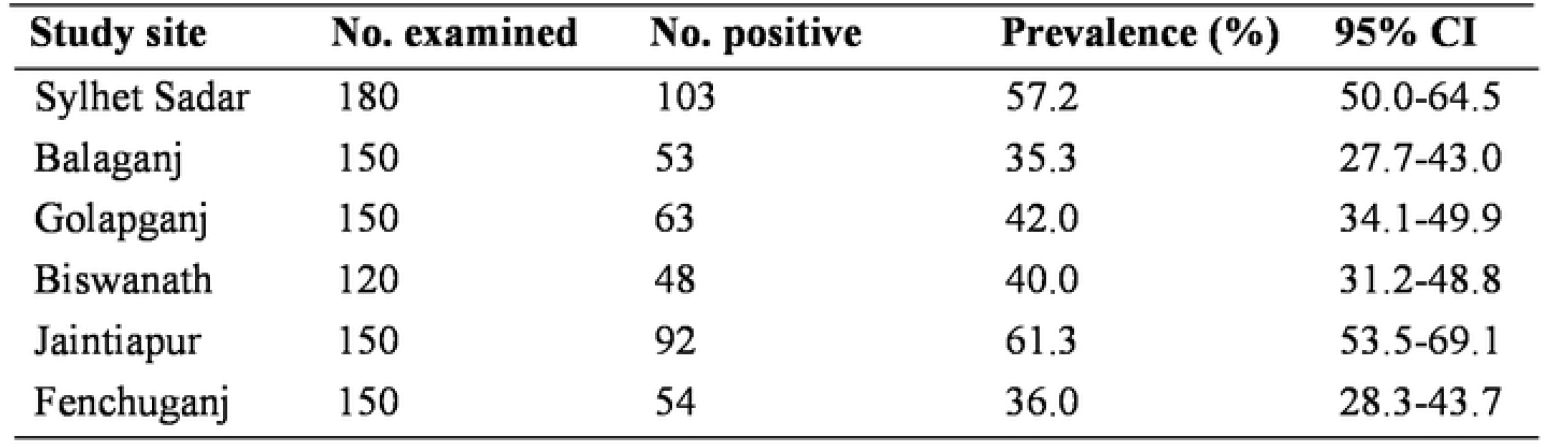
Prevalence of gastrointestinal helminths by study site.

| Study site | No. examined | No. positive | Prevalence (%) | 95% CI |
| --- | --- | --- | --- | --- |
| Sylhet Sadar | 180 | 103 | 57.2 | 50.0-64.5 |
| Balaganj | 150 | 53 | 35.3 | 27.7-43.0 |
| Golapganj | 150 | 63 | 42.0 | 34.1-49.9 |
| Biswanath | 120 | 48 | 40.0 | 31.2-48.8 |
| Jaintiapur | 150 | 92 | 61.3 | 53.5-69.1 |
| Fenchuganj | 150 | 54 | 36.0 | 28.3-43.7 |

**Figure 1.**
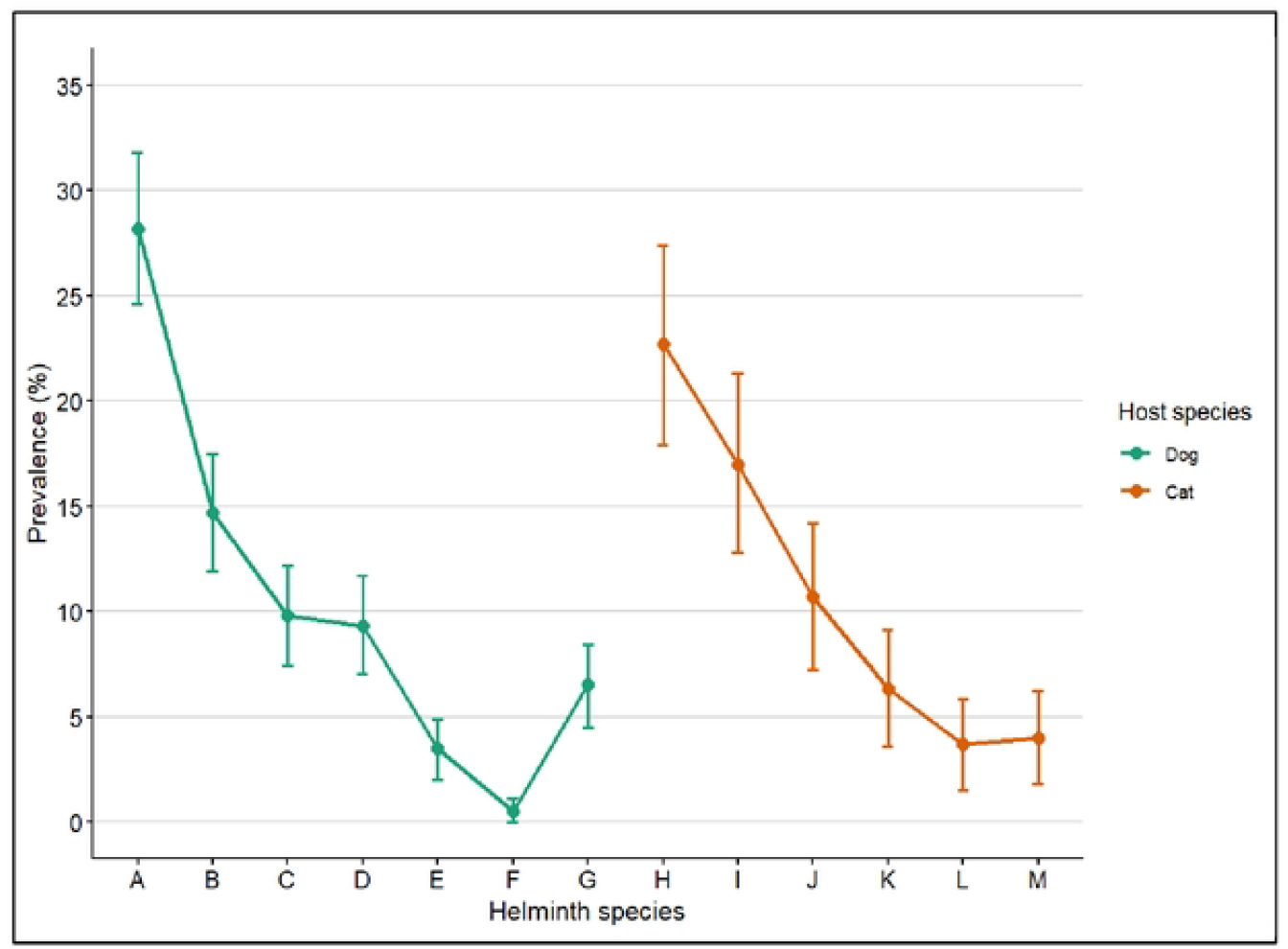
Prevalence of gastrointestinal helminth of dogs and cats in Sylhet. Here, A = Hookworms, B = *Toxocara canis*, C = *Trichuris vulpis*, D = *Dipylidium caninum*, E = *Taenia/Echinococcus* spp., F = *Diphyllobothrium* spp., G = Larvae (unspecified), H = *Toxocara cati*, I = *Ancylostoma tubaeforme*, J = *Hymenolepis* spp., K = *Taenia/Echinococcus* spp., L = *Physaloptera* spp., M = Larvae (unspecified).

**Figure 2.**
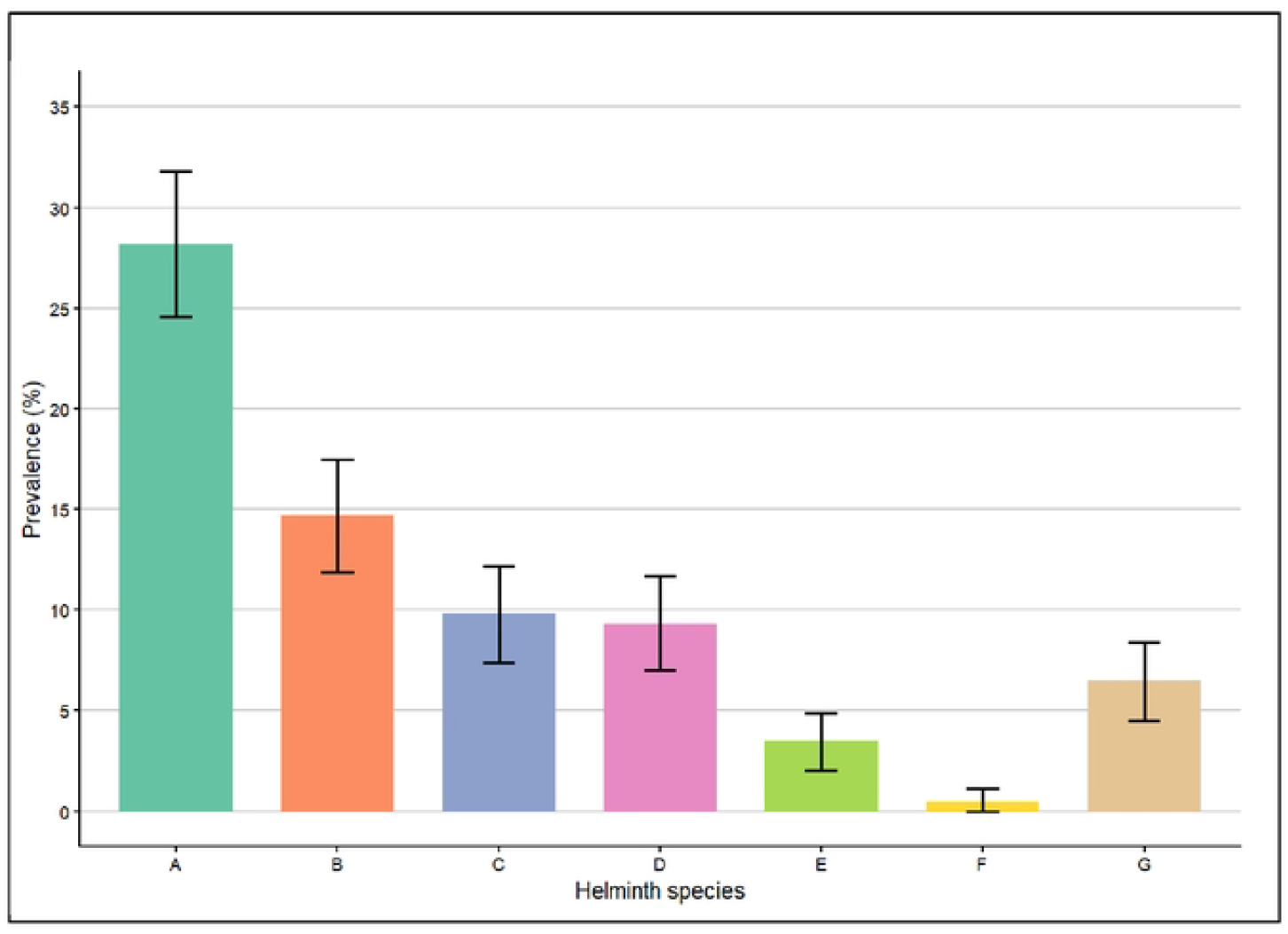
Prevalence of gastrointestinal helminth infection of dogs in Sylhet. Here, A = Hookworms, B = *Toxocara canis*, C = *Trichuris vulpis*, D = *Dipylidium caninum*, E = *Taenia/Echinococcus* spp., F = *Diphyllobothrium* spp., G = Larvae (unspecified).

**Figure 3.**
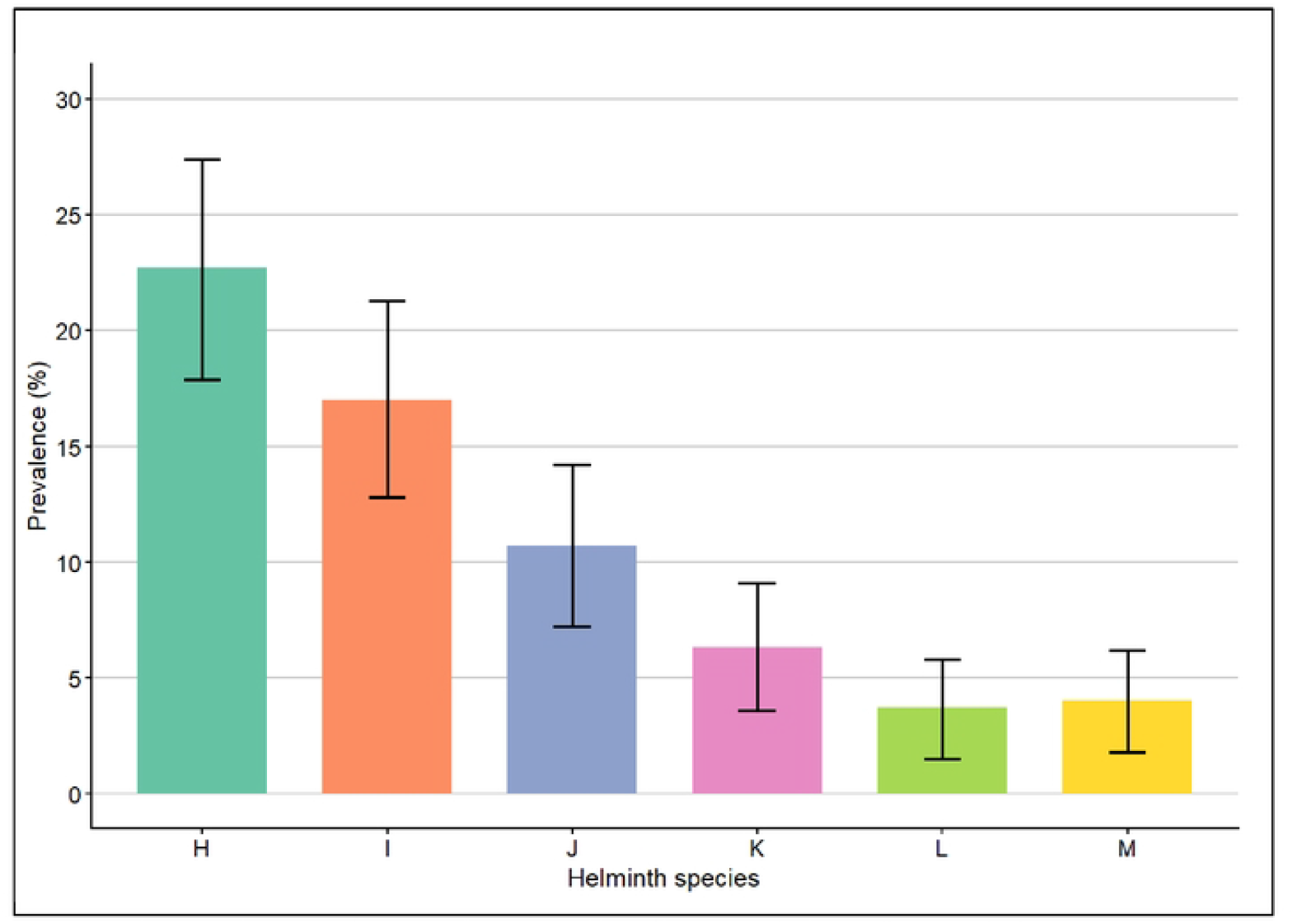
Prevalence of gastrointestinal helminth infection of cats in Sylhet. Here, H = *cara cati*, I = *Ancylostoma tubaeforme*, J = *Hymenolepis* spp., K = *Taenia/Echinococcus* spp., L = *Physaloptera* spp., M = Larvae (unspecified)

**Figure 4.**
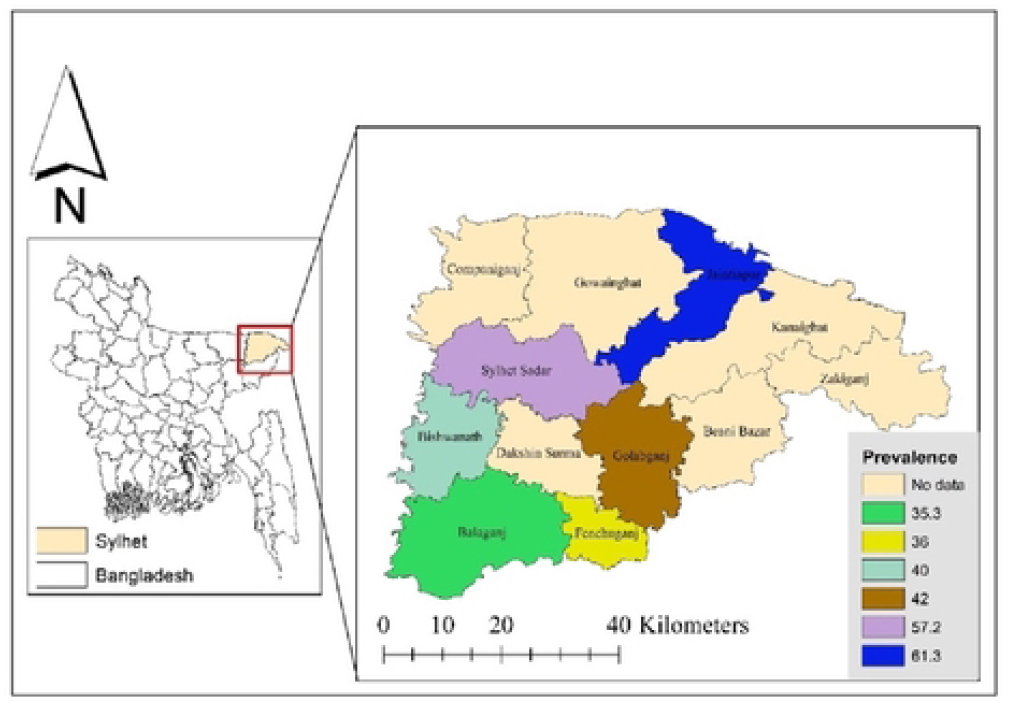
Sub-district-wise prevalence of gastrointestinal helminth infections of dogs and 17 cats in Sylhet district, Bangladesh.

### Overall prevalence of gastrointestinal helminths in dogs

Among the 600 dogs examined, 274 were infected with one or more gastrointestinal helminths, resulting in a prevalence of 45.7%. Hookworms were the most frequently detected parasites (28.2%), followed by *Toxocara canis* (14.7%) and *Trichuris vulpis* (9.8%). Cestode infections, including *Dipylidium caninum* and Taenia/*Echinococcus* spp., were also identified. Mixed infections were recorded in 19.3% (53/274) of infected dogs.

### Overall prevalence of gastrointestinal helminths in cats

Of the 300 cats examined, 139 were positive for gastrointestinal helminths, giving an overall prevalence of 46.3%. *Toxocara cati* was the predominant parasite (22.7%), followed by *Ancylostoma tubaeforme* (17.0%). Cestodes and spirurid nematodes were also detected. Mixed infections were observed in 16.5% (23/139) of infected cats.

### Overall prevalence of gastrointestinal helminths in pet-owned dogs and cats

Among the 600 pet-owned animals (400 dogs and 200 cats), 220 were infected with at least one helminth species, resulting in a combined prevalence of 36.6%. The prevalence was 35.5% in dogs and 39.0% in cats.

### Overall prevalence of gastrointestinal helminths in stray dogs and cats

Stray animals exhibited a substantially higher prevalence of infection. Of the 300 stray animals examined (200 dogs and 100 cats), 193 were infected, yielding an overall prevalence of 64.3%. The prevalence was 66.0% in dogs and 61.0% in cats.

### Helminth prevalence by study site

Marked spatial variation in helminth prevalence was observed across the six study sites. The highest prevalence was recorded in Jaintiapur (61.3%) and Sylhet Sadar (57.2%), while Balaganj and Fenchuganj exhibited comparatively lower prevalence.

### Helminth detection by diagnostic method

The detection rate varied by diagnostic technique. Sheather’s sugar flotation identified the highest number of positive samples, followed by the Formalin–Ether Concentration Technique and the Modified Baermann method (Table 7, Fig 5).

**Table 7.** Gastrointestinal helminths by diagnostic method.

| Diagnostic method | No. positive | Detection rate (%) | 95% CI |
| --- | --- | --- | --- |
| Formalin–Ether Concentration | 354 | 39.3 | 36.1-42.5 |
| Sheather’s Sugar Flotation | 372 | 41.3 | 38.1-44.5 |
| Modified Baermann | 165 | 18.3 | 15.8-20.8 |

**Table 8.** Multivariable logistic regression analysis of risk factors for helminth infection.

| Risk factor | Adjusted OR | 95% CI | p-value |
| --- | --- | --- | --- |
| Stray status | 2.91 | 2.18–3.88 | <0.001 |
| No deworming history | 2.45 | 1.76–3.42 | <0.001 |
| Age <1 year | 1.89 | 1.30–2.75 | 0.001 |
| Outdoor roaming | 2.11 | 1.46–3.07 | <0.001 |
| Rural residence | 1.31 | 0.95–1.80 | 0.096 |

**Table 9.** PCR-confirmed helminths with amplicon size and sequence similarity.

| Host | Parasite Species | Gene Target | Amplicon Size (bp) | No. confirmed | BLAST Similarity (%) | Closest GenBank |
| --- | --- | --- | --- | --- | --- | --- |
| Dog | *Ancylostoma caninum* | ITS-2 | 380–420 | 31 | 99.1–99.8 | *A. caninum* (KF032990) |
|  | *Toxocara canis* | ITS-2 | 400–450 | 18 | 98.7–99.5 | *T. canis* (KY003073) |
|  | *Trichuris vulpis* | ITS-2 | ~350 | 12 | 98.4–99.2 | *T. vulpis* (MT856594) |
|  | *Dipylidium caninum* | 18S rRNA | ~550 | 16 | 99.0–99.6 | *D. caninum* (AY040543) |
|  | *Taenia/ Echinococcus* spp. | cox1 | ~650 | 7 | 98.2–99.0 | *Taenia* sp. (MH792025) |
|  | *Diphyllbothrium* spp. | cox1 | ~650 | 2 | 98.1–98.7 | *Diphyllbothrium* sp. (JN986973) |
| Cat | *Toxocara cati* | ITS-2 | 400–450 | 19 | 98.9–99.7 | *T. cati* (MH043235) |
|  | *Ancylostoma tubaeforme* | ITS-2 | 380–420 | 15 | 98.6–99.4 | *A. tubaeforme* (KF032992) |
|  | *Hymenolepis diminuta* | 18S rRNA | ~550 | 11 | 99.0–99.8 | *H. diminuta* (AF124482) |
|  | *Taenia/Echinococcus* spp. | cox1 | ~650 | 6 | 98.1–98.9 | *Taenia* sp. (MW301188) |
|  | *Physalopter* spp. | ITS-2 | ~360 | 4 | 98.3–99.1 | *Physalopter* sp. (MK070122) |

**Figure 5.**
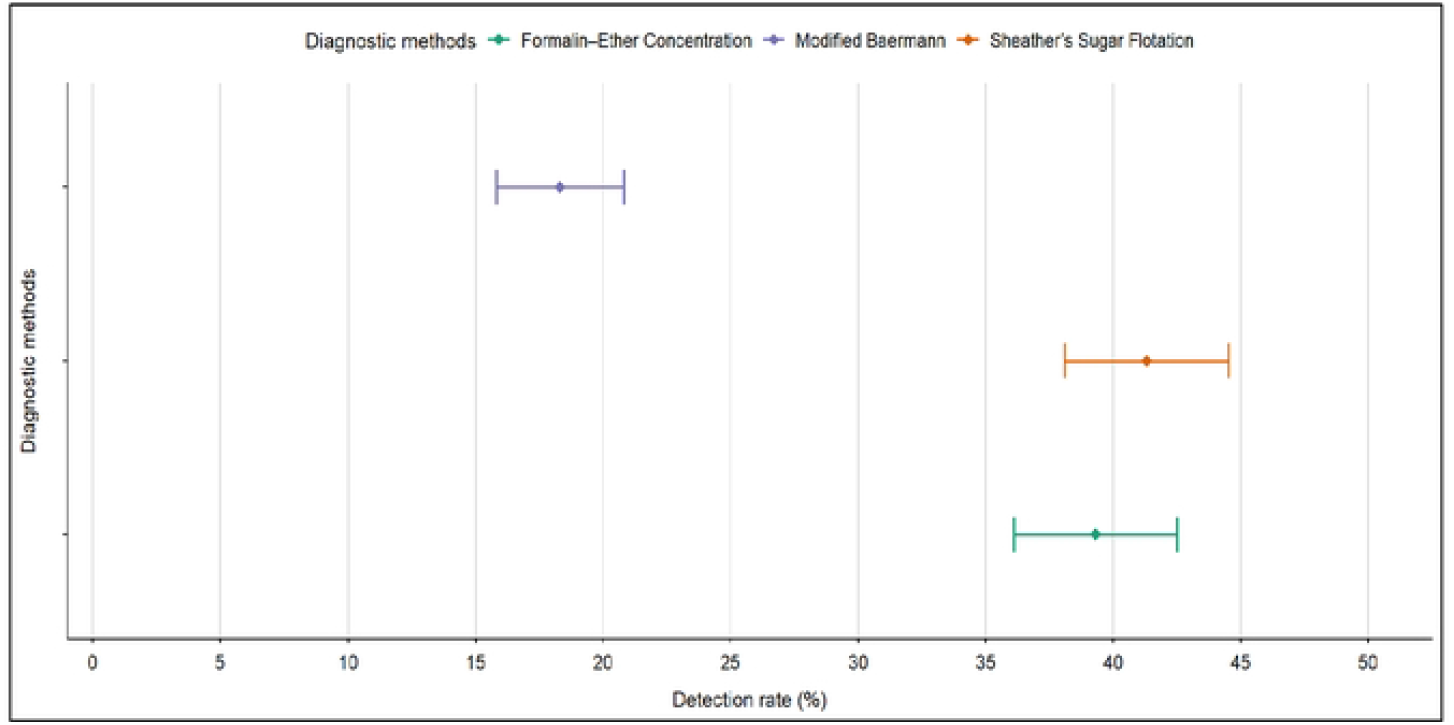
Prevalence detected with various laboratory diagnostic methods

### Risk factors associated with gastrointestinal helminth infection

Multivariable logistic regression analysis demonstrated that stray status, absence of deworming history, age below one year, and outdoor roaming behavior were significantly associated with helminth infection. Rural residence showed increased odds but did not reach statistical significance.

### Molecular Identification and Genomic Validation

Molecular characterization of coproscopically positive fecal samples successfully confirmed parasite identity in 93.0% of cases. Species identification was based on amplicon size, nucleotide sequencing, and comparison with reference sequences deposited in the GenBank database using BLAST analysis. In dogs, *Ancylostoma caninum* was the most frequently confirmed nematode (n=31) using the ITS-2 target, showing a high BLAST similarity of 99.1–99.8% to GenBank accession KF032990. Other canine helminths, including *Toxocara canis* (n=18) and *Trichuris vulpis* (n=12) were validated with similarity scores 98.4%. The use of 18S rRNA and cox1 targets allowed for the precise identification of cestodes such as *Dipylidium caninum* (n=16), *Taenia* spp. (n=7), and *Diphyllobothrium* spp. (n=2). For feline samples, *Toxocara cati* showed the highest confirmation rate (n=19) with 98.9–99.7% similarity to reference strains. Genetic validation of *Ancylostoma tubaeforme* (n=15) and *Hymenolepis diminuta* (n=11) further supported the morphological findings. Notably, *Physaloptera spp*. (n=4) was molecularly confirmed using the ITS-2 region, matching GenBank isolate MK070122 with 98.3–99.1% similarity

## DISCUSSION

This study presents a comprehensive epidemiological and molecular investigation of gastrointestinal helminths in companion dogs and cats in northeastern Bangladesh, providing robust evidence of a substantial parasitic burden and zoonotic potential. The overall prevalence of gastrointestinal helminths (45.9%) highlights the continued circulation of these parasites in companion animal populations, reinforcing concerns that parasitic zoonoses remain underrecognized public health threats in Bangladesh.

The prevalence observed in this study is consistent with reports from other tropical and subtropical regions, where warm temperatures, high humidity, and inadequate sanitation favor the persistence and transmission of helminth eggs and larvae [15]. Comparable prevalence rates have been reported in India, Nepal, and parts of Southeast Asia, indicating that gastrointestinal helminthiasis in companion animals is a regional concern rather than an isolated phenomenon [16,17].

The similar infection rates observed between dogs and cats contrast with findings from high-income countries, where cats often show lower prevalence due to stricter indoor confinement and routine veterinary care [18]. In Bangladesh, however, both dogs and cats frequently roam freely, increasing exposure to contaminated environments and intermediate hosts, which likely explains the comparable prevalence observed.

The predominance of hookworms (*Ancylostoma* spp.) and *Toxocara* spp. in both dogs and cats is of major zoonotic concern. *Ancylostoma caninum* and *A. tubaeforme* are well-documented causes of cutaneous larva migrans in humans, particularly in tropical settings where barefoot exposure is common [19]. Similarly, *Toxocara canis* and *T. cati* are responsible for visceral and ocular larva migrans, conditions that disproportionately affect children and are frequently underdiagnosed in low-resource settings [6].

The detection of *Trichuris vulpis* in dogs, although less prevalent, is epidemiologically relevant, as whipworm infections contribute to chronic gastrointestinal morbidity in animals and may act as indicators of poor environmental hygiene [20]. The identification of cestodes like *Dipylidium, Taenia, Echinococcus, Hymenolepis*, and *Diphyllobothrium* further underscores the complexity of parasite transmission cycles in Bangladesh. These parasites require intermediate hosts, reflecting ecological conditions that support sustained transmission and increasing the risk of human exposure through accidental ingestion of infective stages [7].

The significantly higher prevalence of helminths among stray animals (65.7%) compared with pet-owned animals (36.0%) highlights the critical role of veterinary care and animal management in parasite control. Stray dogs and cats in Bangladesh typically lack access to routine deworming and veterinary supervision, roam extensively, and scavenge for food, all of which increase their exposure to infective stages [21].

The finding that all examined stray cats were infected is particularly concerning and suggests that free-roaming cats may act as major reservoirs for environmental contamination. Similar observations have been reported in urban slum settings in other South Asian countries, where stray animals contribute disproportionately to zoonotic parasite transmission [22]. These findings emphasize the need for integrated stray animal management programs as part of zoonotic disease control strategies.

The marked variation in helminth prevalence across study sites reflects the influence of local environmental and socio-economic conditions. Higher prevalence in Jaintiapur and Sylhet Sadar may be associated with higher animal density, frequent human-animal interactions, and inadequate waste management. Environmental contamination of soil and water has been identified as a key driver of helminth transmission in similar ecological settings [23]. Such spatial heterogeneity highlights the importance of localized epidemiological data for designing targeted interventions. Blanket national strategies may fail to address site-specific drivers of infection, underscoring the value of region-specific surveillance in Bangladesh.

The use of multiple coproscopic techniques improved parasite detection, with Sheather’s sugar flotation demonstrating the highest detection rate. This finding aligns with previous studies reporting superior sensitivity of flotation techniques for detecting light eggs such as those of hookworms and *Toxocara* spp. [24]. The Modified Baermann technique proved essential for detecting larval stages, reinforcing the need for complementary diagnostic approaches in epidemiological surveys.

Multivariate analysis identified stray status, absence of deworming, young age, and outdoor roaming as significant risk factors for helminth infection. These findings are consistent with studies conducted in other endemic regions, where management practices and behavioral factors play a more decisive role than geographic location alone [25]. The increased susceptibility of young animals likely reflects immature immune responses and higher exposure during exploratory behaviour.

The high PCR confirmation rate (93%) in this study validates the reliability of traditional coproscopic techniques while demonstrating the indispensable value of molecular tools for species-level differentiation. The high nucleotide similarity observed between the Sylhet isolates and global zoonotic reference strains suggests a degree of genetic homogeneity and indicates potential regional or international transmission networks. Specifically, the molecular confirmation of *Taenia* and *Hymenolepis* is of critical public health importance, as these parasites are linked to severe human morbidity [16, 26]. The detection of *Ancylostoma caninum* and *A. tubaeforme*, both confirmed with high sequence fidelity, reinforces the risk of cutaneous larva migrans in the local population, particularly where environmental hygiene is suboptimal [8]. These findings advocate for the routine integration of molecular diagnostics into national One Health surveillance to accurately assess zoonotic risks in Bangladesh.

The findings of this study underscore the interconnectedness of animal health, human health, and environmental factors in Bangladesh. Companion animals, particularly stray dogs and cats, serve as important reservoirs of zoonotic helminths, contributing to environmental contamination and human exposure [18]. The high prevalence of zoonotic species calls for integrated One Health interventions that combine veterinary services, public health education, environmental sanitation, and policy support. Regular deworming of companion animals, improved waste management, public awareness campaigns on hygiene, and coordinated stray animal control programs are essential components of an effective One Health strategy. Without such integrated efforts, gastrointestinal helminths will continue to pose a silent but significant threat to both animal welfare and human health in Bangladesh.

This study provides valuable baseline information while also highlighting opportunities for further research. The cross-sectional design offers an important snapshot of helminth infections in the study population, though longitudinal approaches could strengthen causal interpretations in future work. Additionally, incorporating quantitative egg counts would enhance understanding of infection intensity. Expanding future studies to include larger sample sizes and a wider range of host species would further enrich knowledge of the epidemiology of helminth infections in the region.

## CONCLUSIONS

The high prevalence of zoonotic gastrointestinal helminths in Sylhet underscores the urgent need for a multi-sectoral One Health approach to mitigate public health risks in northeastern Bangladesh. Given that stray status was identified as the most significant risk factor, priority must be given to integrated stray animal management, including mass deworming programs and the establishment of community-led veterinary surveillance. Public health authorities should implement targeted education campaigns focusing on hand hygiene and the risks associated with environmental contamination, particularly in high-prevalence zones. Furthermore, the molecular evidence of genetic similarity between local isolates and global zoonotic strains necessitates the creation of a national parasitic sequence database to track transmission dynamics. Integrating veterinary diagnostics with pediatric clinical screenings will facilitate earlier diagnosis and reduce the long-term morbidity associated with these neglected tropical diseases in vulnerable populations.

## COMPETING INTERESTS

There is no conflict of interest declared by any of the authors.

## ACKNOWLEDGMENTS

This work was supported by the Sylhet Agricultural University Research System. The authors wish to acknowledge the support of the KDCA-DVP 2024 Grant Program, Division of Vectors & Parasitic Diseases, Korea Disease Control and Prevention Agency (KDCA). SAURES and KDCA had no role in the study design, data collection, and analysis, decision to publish, or preparation of the manuscript

## AVAILABILITY OF DATA AND MATERIALS

The datasets and resources used in this current study are available from the corresponding author upon reasonable request.

## ETHICAL APPROVAL

This study protocol was reviewed and approved by the Sylhet Agricultural University Research System (SAURES-UGC-2023-2024-02) and the Department of Parasitology, Sylhet Agricultural University, Bangladesh.

## AUTHOR CONTRIBUTIONS

All authors reviewed and provided feedback for this manuscript. The final version of this manuscript was vetted and approved by all authors.

## Notes

### Competing Interest Statement

The authors have declared no competing interest.

